# Nanoarchaeosome-mediated epirubicin delivery induces sustained intracellular stress and suppresses adaptive glioblastoma phentoypes

**DOI:** 10.64898/2026.07.06.736470

**Authors:** Adhithya Subramanian Gopalakrishnan, Subastri Ariraman, Sourav Ganguli, Adithya Hitesh, Shahid Mohammad, Mukilarasi B, Swathi Sudhakar, Pavithra L Chavali

## Abstract

Although anthracyclines such as epirubicin are potent DNA-damaging agents, their application in glioblastoma (GBM) is limited by poor intracellular penetration, lack of durable responses and rapid emergence of adaptive tumor phenotypes. Here, we demonstrate that nanoarchaeosome-mediated delivery of epirubicin (NanoEpi) enables functional reprogramming of GBM survival under therapeutic stress. Nanoarchaeosomes composed of archaeal ether lipids exhibited high encapsulation efficiency (∼96%) and nanoscale stability. Although both Epi and NanoEpi showed similar bulk uptake, both in established (U251-MG) and patient-derived (Gli5) glioblastoma models, NanoEpi induced significantly greater cytotoxicity than free epirubicin, indicating enhanced intracellular drug engagement. NanoEpi induced enhanced DNA damage, elevated reactive oxygen species, and mitochondrial depolarisation, leading to cytoskeletal collapse. In 3D gliomasphere systems, NanoEpi showed improved penetration and sustained retention, resulting in suppression of core viability and invasion. This correlates with its increased uptake by the lysosomes. Notably, even a transient exposure led to depletion of sphere-initiating capacity and complete loss of clonogenic recovery, indicating targeting of the stem-like compartment. This was accompanied by attenuation of MMP-2/9 activity and reduced angiogenic signalling in a chorioallantoic membrane model. These findings establish nanoarchaeosomes as a robust lysosome-directed delivery platform that extends beyond passive drug transport to sustain intracellular stress and suppress invasive adaptation and limit recurrence in GBM.

## 1. Introduction

Glioblastoma (GBM) is one of the most aggressive and treatment-refractory malignancies, characterised by profound intratumoral heterogeneity, diffuse invasion, and a highly adaptive cellular hierarchy that enables rapid therapeutic escape. A defining feature of GBM progression is the persistence of stem-like tumor cell populations that survive cytotoxic stress, reinitiate tumor growth, and drive recurrence [1]. Consequently, effective therapeutic strategies must extend beyond bulk cytotoxicity to disrupt the functional resilience and regenerative capacity of these tumor-propagating compartments [2].

Epirubicin (Epi), a semi-synthetic anthracycline and 4′-epimer of doxorubicin, is a potent DNA-intercalating chemotherapeutic with less cardiotoxicity and myelosuppression than conventional anthracyclines [3]. Epirubicin is efficacious in various solid tumors, but its use in glioblastoma (GBM) is limited by poor blood-brain barrier penetration, low tumor retention and limited intracellular persistence [4–6]. To address these challenges, alternative delivery strategies, such as convection-enhanced delivery and nanoparticle-based formulations, have been investigated to enhance brain tissue penetration and therapeutic concentrations [2, 7, 8] Additionally, liposomal formulations of epirubicin, including PEGylated and non-PEGylated forms, have demonstrated reduced cardiac dysfunction while maintaining comparable antitumor efficacy [9].

Nanoarchaeosomes present a next-generation strategy to overcome delivery limitations [10–13]. Nanoarchaeosomes, composed of archaeal ether-linked phospholipids, exhibit superior structural stability, resistance to oxidative stress, and reduced susceptibility to enzymatic degradation compared to conventional liposomes [14–16]. These properties can enhance drug encapsulation efficiency, prolong systemic circulation, and improve stability under physiological conditions and acidic tumor microenvironment. Their membrane characteristics allow surface functionalization for blood-brain barrier targeting, improved membrane fusion, and controlled drug release. The intrinsic biocompatibility and lower immunogenicity of nanoarchaeosomes further support their suitability as carriers for anthracyclines such as epirubicin.

Here, we investigated whether nanoarchaeosome-mediated epirubicin delivery can extend beyond passive enhancement of drug uptake to actively suppress adaptive tumor survival programs in GBM. Specifically, we investigate whether nanoarchaeosome-loaded epirubicin (NanoEpi) can alter the functional trajectory of GBM cells by disrupting survival pathways associated with stemness, invasion, and post-treatment recovery. Using a combination of 2D cultures, patient-derived 3D gliomasphere models, and a vascularized chorioallantoic membrane (CAM) system, we demonstrate that NanoEpi not only enhances cytotoxic efficacy but also induces a sustained suppression of tumor cell plasticity. This could be due to the accumulation of NanoEpi within the lysosomal compartments leading to induction of sustained stress signalling. As a consequence, even a transient exposure to NanoEpi can substantially reduce sphere-initiating capacity (tumor adaptation), inhibition of matrix invasion, attenuation of angiogenic signalling, and prevention of proliferative rebound following drug withdrawal.

## 2. Materials and Methods

### Synthesis and Characterization of NanoEpi

Nanoarchaeosomes (Nanoarch) were prepared as previously described [14]. These carriers were created by incorporating 20% archaeal lipids extracted from *Aeropyrum pernix* K1 and 80% 1-stearoyl-2-oleoyl-sn-glycero-3-phosphocholine (SOPC, Sigma) in a weight/volume (w/v) ratio to achieve a final concentration of 1 mg/ml. The nanovesicles were formed by hydration in Milli-Q water and sonication, and were then used to load epirubicin. Epirubicin-loaded nanoarchaeosomes (NanoEpi) were prepared by mixing equal volumes of epirubicin (Epi) and Nanoarch to reach a final concentration of 200 µM Epi. This mixture was stirred at 300 rpm for 12 hours at room temperature. To facilitate the efficient encapsulation of Epi in the Nanoarch, the mixture underwent multiple freeze-thaw cycles. The NanoEpi was filtered through a 30 kDa Amicon ultra-centrifugal filter for 30 minutes at 13,000 rpm. The concentration of free drug in the filtrate was measured at 565 nm using an epirubicin standard curve to estimate the encapsulation efficiency.

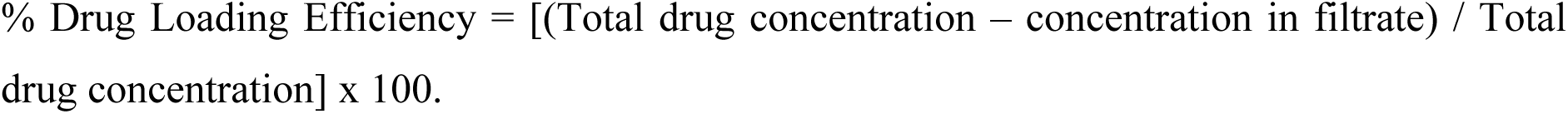

The Nanoarch and NanoEpi were characterized using scanning electron microscopy (S-4800 Hitachi) to assess size distribution by acquiring images at 1-5 kV after gold sputtering (Polaron SC7640). Fourier Transform Infrared (FTIR) profiles of Nanoarch, Epi, and NanoEpi were obtained to confirm Epi loading into Nanoarch, generating a transmittance spectrum in the Nicolet iS5 FT-IR over the range of 500-4000 cm^-1^ after pelleting in a 1:100 KBr (% w/w) mixture.

### Cell Culture

Gli5 is a primary cell culture established from a Grade 4 Glioblastoma tissue obtained from a 52-year-old male patient who had undergone surgical resection. The tissue was collected after approval of the institutional ethical committee (IISERTPT/2024/002 & EC/NIMS/3453/2024, NIMS, Hyderabad) and with the patient’s informed consent. This cell line is highly proliferative and retains wild-type p53 and ATRX. The tissue obtained after surgery was minced and dissociated using a 0.5% trypsin-EDTA solution along with collagenase. The cells were filtered through a cell strainer and cultured in DMEM Glutamax containing 20% FBS and 1x penicillin-streptomycin (Gibco). The U251-MG cells (ATCC) used in this study were cultured in DMEM Glutamax supplemented with 10% FBS and 1x penicillin-streptomycin (Gibco). All cultures were incubated at 37°C under 5% CO_2_ in a humidified chamber. Mycoplasma contamination was tested and found to be negative by PCR. All experiments were conducted with Gli5, unless otherwise specified. For tumor spheroids, cells were grown in 96-well U-bottom ultra-low attachment plates (174925, Nunclon Sphera, Thermo) in DMEM-F12 media supplemented with N2, B27 (Gibco), and 20 ng/ml of EGF and bFGF (V-Edit). Day 5 spheroids were used for experiments.

### MTT Assay for Cell Viability

Epi and NanoEpi at indicated concentrations were serially diluted in media and applied to 3000 cells, which had been allowed to attach and grow overnight. After 24 hours of drug treatment, the media was removed, and 0.5 mg/ml of MTT (475989, Sigma) was added. The cells were then incubated in the dark for 4 hours. The formazan crystals were dissolved in 100 µl of DMSO (TC349, Himedia) per well, and absorbance was measured at 570 nm using a microplate reader (BioTek Cytation 5, Agilent).The percentage viability was calculated using the following formula:

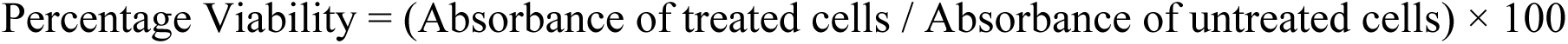

The data was plotted, and the IC50 value was determined using GraphPad Prism by fitting the curve with a non-linear regression model.

### Cell Cycle Analysis

Cells were treated with 5 µM Epi and NanoEpi for 24 hours, and their natural fluorescence from epirubicin was observed using an inverted fluorescence microscope (Nikon Eclipse Ts2). After trypsinization and washing with 1x PBS, cells were fixed in 70% ethanol for 30 minutes at 4°C, and were washed again with PBS. For cell cycle analysis, the cell pellet was stained with 50 µg/ml propidium iodide (P4170, Merck), 0.1% Triton-X 100, and 100 µg/ml DNase-free RNase A (EN0531, Thermo) for 30 minutes at room temperature. Cell cycle profiles were assessed using the CytoFLEX S Flow Cytometer (Beckman Coulter).

### Matrigel Invasion Assay

After drug treatment, the tumor spheroids were embedded in a drop of Matrigel (354234, Corning). After solidification, DMEM Glutamax with 10% FBS was overlayed for a gradient chemoattraction. Invasion was observed and captured daily using a brightfield microscope at 4x magnification. On day 3, invading tumor spheroids were fixed in 4% PFA (I28800, Thermo) for 20 minutes and stained with 1 µg/ml DAPI (D9542, Merck) and Phalloidin (Actin-555, R37112, Invitrogen) for 30 minutes at room temperature. The invading spheroids were mounted on glass slides using mounting media (Prolong Gold, P36930, Invitrogen) and imaged using a confocal microscope (FV3000, Olympus).

### Immunostaining

The cells/tumor spheroids were fixed with 4% PFA for 20 minutes in room temperature after 24 hours of drug treatment. The samples were permeabilised with 0.1% TritonX-100 for 20 minutes at room temperature after a PBS wash. The cells/spheroids were blocked with 5% BSA (Himedia) in PBS with 0.1% Tween20 for 1 hour. The primary antibody was diluted in the blocking solution and incubated with the samples overnight at 4 °C. PBS with 0.1% Tween20 was used as the washing buffer. After 3 washes of 5 minutes each, the samples were incubated with the secondary antibody conjugated to an appropriate fluorophore diluted in the blocking solution for 2 hours at room temperature. After another 3 washes of 5 minutes each, 1 µg/ml DAPI was added and incubated for 10 minutes. The samples were washed once in PBS before mounting on a glass slide with antifade mountant and after mounting, the slides were stored at 4°C before image acquisition in the confocal microscope (FV3000, Olympus). The cells/spheroids were also stained with phalloidin Actin555 ready probes (Invitrogen), if mentioned, according to the manufacturer’s protocol.

The primary antibodies used for immunostaining include Phospho-Histone H2A.X (Ser139) (20E3) Rabbit Monoclonal Antibody CST-9718 (1:500), Cleaved Caspase-3 (Asp175) Antibody CST-9661 (1:500) and Ki-67 (8D5) Mouse Monoclonal Antibody CST-9449 (1:1000). The anti-rabbit and anti-mouse IgG (H+L) highly cross-adsorbed secondary antibodies with Alexa Fluor 488 (Invitrogen) were used in a 1:1000 dilution.

### Staining for Lysosomes, ROS and Mitochondrial membrane potential

After 24 hours of Epi and NanoEpi treatment, cells were stained with LysoTracker Green DND-26 (75 nM) in DMEM for 30 minutes at 37°C. They were then treated with Hoechst (0.5 µg/ml) for 20 minutes, washed with PBS, and imaged live using a confocal microscope at 60x magnification.

For DCFDA and JC-1 staining, cells were treated with the reagents 24 hours after NanoEpi exposure. Intracellular ROS levels were assessed by incubating cells with 20 μM DCFDA for 30 minutes at 37°C, using calibration controls for fluorescence validation. Mitochondrial membrane potential was evaluated with 1 μg/ml JC-1, also for 30 minutes at 37°C, with appropriate controls for specificity. Fluorescence images were captured using a Nikon Eclipse Ti-U microscope following PBS washes.

### Colony formation assay

After 24 hours of drug treatment, the cells were washed in PBS twice and trypsinized. A total of 500 viable cells from all categories were seeded into each well of a 6-well cell culture plate. The plate was left undisturbed for 10 days, with partial media changes every third day. Once colony formation was visible, the cells were fixed with ice-cold methanol for 10 minutes and stained with 0.5% crystal violet (V5265, Merck) for 30 minutes at room temperature. The plates were washed thrice with PBS and imaged in a Gel documentation apparatus (GelDoc Go, Bio-Rad)

### Extreme Limiting Dilution Assay (ELDA)

Following 24 h of drug treatment, the cell suspension was serially diluted in tumor spheroid medium to obtain densities ranging from 10 to 1000 cells per ml. Subsequently, 100 μl of the diluted suspension was seeded into each well of a 96-well ultra-low attachment plate. Plates were centrifuged briefly to facilitate cell settling, and spheroid formation was monitored over time. Wells were scored as positive or negative for spheroid formation based on predefined morphological criteria. Fresh medium was added every 3 days, and after 10 days, wells were evaluated for the presence or absence of spheroids. ELDA analysis was then performed using the number of positive wells relative to the total wells seeded, and the corresponding graph was generated using the ELDA calculator [17].

### Western Blotting

The conditioned media from the spheroids collected at the time point of 48 hours of recovery after 24 hours of drug treatment were analysed for MMP levels. The collected supernatant was concentrated using a SpeedVac (Martin Christ) and loaded onto a 10% polyacrylamide gel under denaturing conditions. The samples were resolved and transferred onto a nitrocellulose membrane. The membrane was blocked in 5% non-fat dry milk in TBS with 0.1% Tween 20. The primary antibody diluted in the blocking solution was probed overnight at 4°C, and the HRP-conjugated anti-rabbit IgG secondary antibody was incubated for 1 hour at room temperature with constant rocking. The membrane was washed 3 times with TBS containing 0.1% Tween 20 after both primary and secondary antibody incubations. The blots were imaged using chemiluminescence (Clarity, Biorad) in an Amersham ImageQuant 800 imager (Cytiva).The antibodies used includes MMP-2 (D4M2N) Rabbit Monoclonal Antibody CST-40994 (1:1000) and HRP linked anti-rabbit IgG CST-7074 (1:5000)

### Gelatin Zymography

Equal volumes of supernatant from all treatments were resolved on an 8% polyacrylamide gel containing 0.25 mg/ml Gelatin (MB169, Himedia) and SDS under non-reducing conditions. Once samples resolved, the gel was washed in wash buffer containing 2.5% TritonX 100, 50 mM Tris-HCl, pH 7.6, 5 mM CaCl_2_ and 1 µM ZnCl_2_ for 1 hour. Subsequently, the gel was incubated in buffer containing 1% TritonX 100, 50 mM Tris-HCl, pH 7.6, 5 mM CaCl_2_ and 1 µM ZnCl_2_ for 16 hours at 37°C. Later, the gel was stained with 0.5% Coomassie Blue R250 in 40% methanol and 10% acetic acid. After destaining, the digested bands were imaged in a gel documentation system (Bio-Rad).

### Chorioallantoic membrane assay and quantitative PCR

The chorioallantoic membrane assay was performed to assess the anti-angiogenic effects of the NanoEpi treatment, as described previously [15]. Briefly, we established a tumor microenvironment on fertilised day 5 growing chick embryos (*Gallus domesticus* L.) incubated in a 37°C chamber with around 60% relative humidity by embedding a coverslip with Gli5 cells. 3000 cells were seeded onto coverslips, and 4 µM drug was applied locally. We quantified morphological changes in blood vessels in the basal, tumour-induced, and tumor+ NanoEpi-treated categories by capturing the CAM vasculature with a stereomicroscope (Olympus) at different time points. The images were evaluated using the IKOSA CAM tool to obtain vessel area, length, and the number of nodes.

The gene expression levels of angiogenic markers, including vascular endothelial growth factor A (VEGFA), fibroblast growth factor 2 (FGF2), and angiopoietin 1 (ANG1), were determined in CAM vasculature regions within different treatment regimens using quantitative real-time PCR. RNA from the excised tissue regions were isolated using Trizol (Invitrogen), and cDNA was synthesised. SYBR FAST qPCR Kit (Kapa Biosystems) was used for the qPCR reaction set in the Eppendorf Realplex 4 Mastercycler system according to the manufacturer’s instructions. Chicken β-actin levels were used as an internal control to normalise relative gene expression levels quantified using the 2^-ΔΔCt^ method.

### Image Analysis

Image analysis of drug uptake, immunostaining and invasion assays was done using ImageJ. The mean fluorescence intensity of epirubicin uptake was estimated after background subtraction, normalised per tumour sphere. The relative fluorescence intensity of pH2AX was calculated by normalising with DAPI intensity. The area of Matrigel invasion was quantified by manually subtracting the core and background regions using thresholding. The data points for the line profile of individual channels to show colocalization were generated in ImageJ.

### Statistical analysis

All data are represented in Mean ± SD from at least 3 biological replicates. For most assays, a One-Way ANOVA followed by an appropriate multiple-comparisons test (Dunnett’s multiple comparison) was performed to determine significance. For the CAM assay, we performed a two-way ANOVA followed by Tukey’s multiple-comparison test in GraphPad Prism 10.5.0. A P-value of <0.05 is considered statistically significant.

## 3. Results and discussion

### 3.1. Nanoarchaeosome formulation and characterization

Nanoarch and NanoEpi produced uniform, non-aggregated spherical vesicles with median diameters of ∼42 nm (Nanoarch) and ∼30 nm (NanoEpi), respectively (**Figure 1A**). The data indicate a consistent nanometer-scale morphology suitable for drug delivery. To confirm that epirubicin was structurally retained in NanoEpi post-loading and purification, FTIR spectroscopy was performed. Ether-stretch bands, archaeal lipid fingerprints, and characteristic anthracycline aromatic peaks were preserved in NanoEpi, confirming co-retention of carrier and cargo rather than non-specific adsorption (**Figure 1B**). Encapsulation efficiency was quantified without dye conjugation by measuring intrinsic fluorescence of epirubicin. Filtrate absorbance was read at 565 nm and interpolated to epirubicin standard curve yielding 96.16 ± 1.8% encapsulation of the drug out of total loaded.

**Figure 1.**
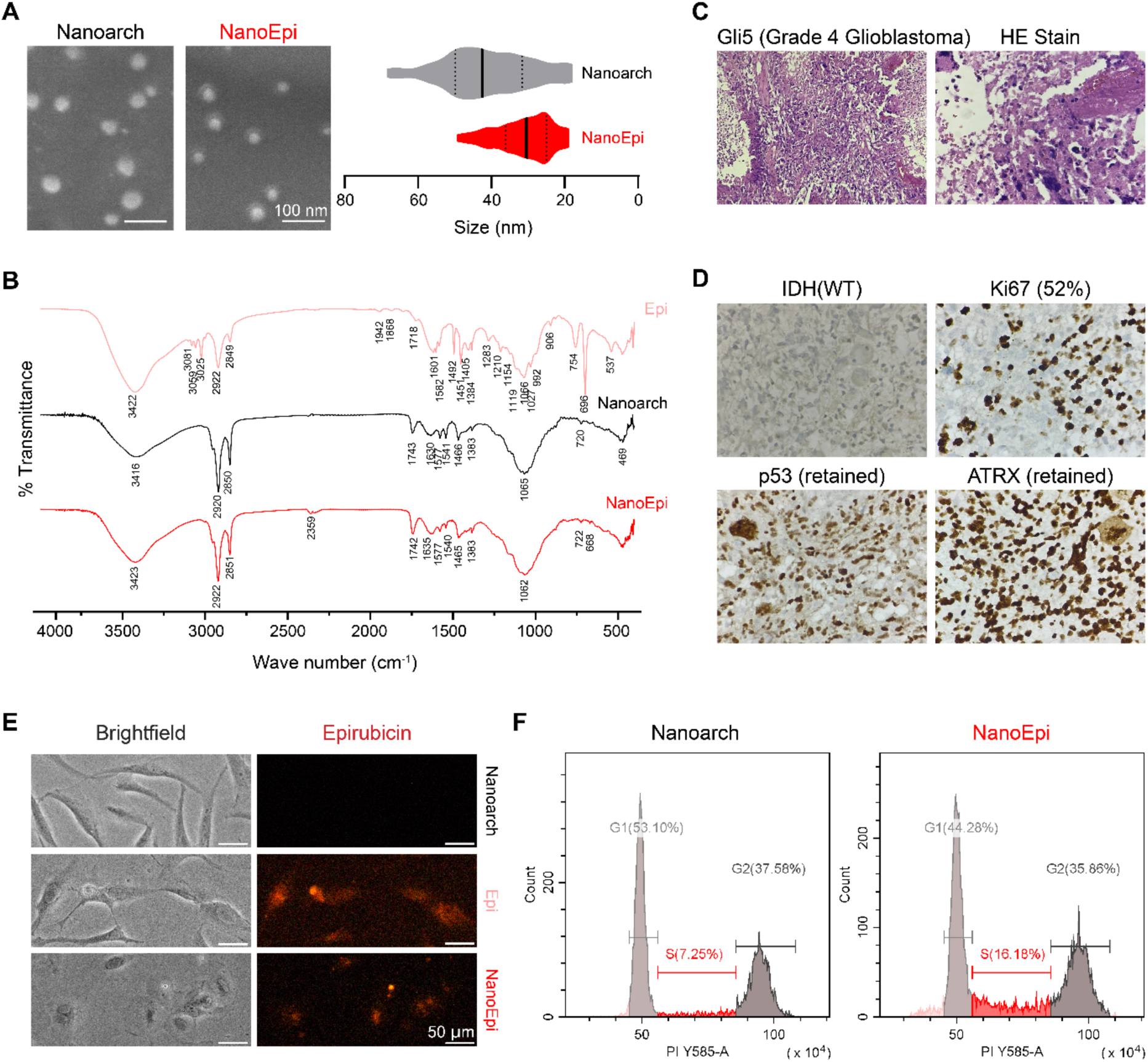
Molecular characterization and Cellular Uptake of Nanoarchaeosome-Loaded Epirubicin. **(A)** Representative scanning electron microscopy (SEM) images and corresponding size distribution analysis (violin plots) for free nanoarchaeosomes (Nanoarch) and epirubicin-loaded nanoarchaeosomes (NanoEpi). **(B)** Fourier-transform infrared (FTIR) spectroscopy spectra comparing free Epirubicin (Epi), free nanoarchaeosomes (Nanoarch), and the nano-formulation (NanoEpi) to confirm chemical integration and stability. **(C)** Histopathological analysis of Patient-Derived Gli5 (Grade 4 Glioblastoma) tissue using Haematoxylin and Eosin (HE) staining. **(D)** Immunohistochemical (IHC) profiling of the GBM sample, showing IDH (wild-type) status, Ki67 proliferative index (52%), and retention of p53 and ATRX markers. **(E)** Brightfield and fluorescence microscopy images demonstrating cellular uptake in glioblastoma cells. Red fluorescence indicates the localization of Epirubicin (Epi) and Nanoarchaeosome-loaded Epirubicin (NanoEpi) compared to the non-fluorescent Nanoarch control. **(F)** Flow cytometry histograms for Nanoarch and NanoEpi treatments showing the distribution of cell populations across different cell cycle phases (G1, S and G2).

Gli5 is a male patient-derived primary Glioblastoma cell line with wild-type p53, IDH and retained ATRX (**Figure 1C-1D**). Additionally, we used standard U251-MG cells which is a high-grade astrocytoma cell line with mutant p53. We first treated the cells with 5 µM free Epi or NanoEpi for 24 hours. Drug distribution was tracked by fluorescence microscopy using epirubicin autofluorescence, revealing that gross uptake intensities were similar between Epi and NanoEpi after quantification (**Figure S1A**). Additionally, NanoEpi induced ∼50% more cells in S-phase by cell-cycle cytometry (5 µM, 24 h) relative to Nanoarch control (7.25% in S-phase vs 16.18% cells in S-phase for NanoEpi), supporting functional proliferative arrest rather than cytostatic survival (**Figure 1F**). A similar S-phase arrest was also observed in Epi-treated cells, as reported [18] (**Figure S1B**), suggesting that NanoEpi functionally reproduces the effects of Epirubicin, resulting in significant S-phase arrest.

### 3.2 NanoEpi enhances intracellular stress responses in GBM cells

We next compared the viability of cells treated with 1-5 μM Epi or NanoEpi and observed that both the NanoEpi-treated Gli5 and U251-MG cell lines responded better, wherein we found ∼50% Gli5 cells were non-viable at 5 μM NanoEpi treatment (**Figure 2A**). We then determined the IC₅₀ values and found that NanoEpi had an IC₅₀ of 2.8 μM in Gli5 cells (**Figure 2B**). In contrast, epirubicin treatment did not induce significant cell death even at higher concentrations, making it difficult to determine an IC₅₀. These results were also verified in U251-MG cells where the IC₅₀ of NanoEpi was 4.4 μM (**Figure S2A**). Since epirubicin is known to induce DNA damage, we investigated genotoxic stress induced by Epi and NanoEpi treatment, respectively. We checked the live/dead cell ratio after Epi and NanoEpi treatment using Calcein AM and PI staining, which also showed a similar trend, with NanoEpi increasing cell death in Gli5 cells (**Figure S2B**). Despite similar uptake, NanoEpi showed an increase in nuclear DNA damage, as visualised by the anti-γH2AX-Ser139 antibody (**Figure 2C**). Additionally, NanoEpi induced significantly higher γH2AX fluorescence intensity in Gli5 cells, while the p-H2AX-positive cell fraction remained elevated compared to free epirubicin (**Figure 2D**). This suggests that NanoEpi increases the DNA damage burden per cell, in addition to increasing the proportion of responsive cells. This distinction cannot be made solely from uptake measurements. Under cytotoxic stress, many chemotherapeutics trigger actin cytoskeleton remodeling as part of cell-death signaling and a stress-adaptation response [19]. Cytoskeletal stress phenotypes were co-visualised by phalloidin revealing NanoEpi-specific actin collapse and cellular dysmorphology (**Figure 2D**). NanoEpi treatment also displayed an elevated ROS, as measured by DCFDA staining (20 µM, 30 min) (Figure 2E), and reduced mitochondrial membrane potential (Δψm), assessed by JC-1 (5,5′,6,6′-Tetrachloro-1,1′,3,3′-tetraethylbenzimidazolylcarbocyanine iodide) ratiometric imaging, in which depolarised mitochondrial membrane potential stains green due to JC-1 monomers (**Figure 2F**). Together, these findings demonstrate enhanced intracellular stress flux in NanoEpi-treated cells, despite no detectable changes in overall epirubicin uptake.

**Figure 2.**
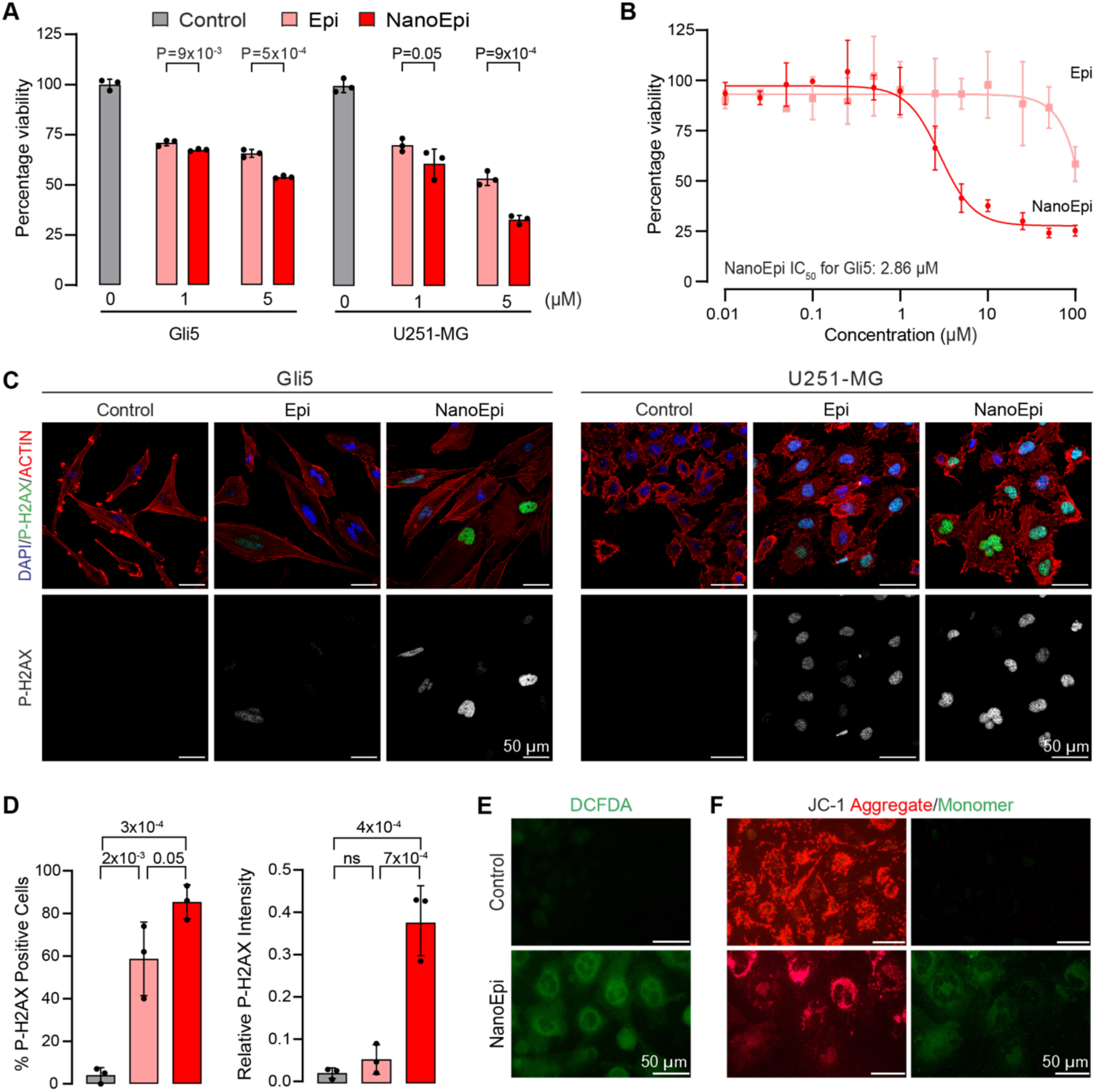
In vitro therapeutic efficacy, DNA damage, and mitochondrial stress in GBM cells upon NanoEpi treatment. **(A)** Comparative cell viability (percentage) of Gli5 and U251-MG glioblastoma cell lines after treatment with free Epi and NanoEpi at 1 μM and 5 μM concentrations. **(B)** Dose-response curves determining the IC_50_ value for NanoEpi in Gli5 cells, measured as 2.865 μM. **(C)** Immunofluorescence imaging of Gli5 and U251-MG cells stained with p-H2AX (green), Actin (red) and DAPI (blue) to visualize drug-induced DNA damage. **(D)** Quantitative analysis of DNA damage in Gli5 cells, showing a significant increase in the percentage of p-H2AX positive cells and relative p-H2AX intensity in NanoEpi-treated cells compared to Epi. **(E)** Reactive oxygen species (ROS) detection in Gli5 cells using DCFDA staining, comparing Control and NanoEpi-treated groups. **(F)** Mitochondrial membrane potential analysis using JC-1 staining; the transition from red aggregates to green monomers indicates mitochondrial depolarization in NanoEpi-treated cells.

### 3.3 NanoEpi improves penetration and suppresses invasion in 3D gliomaspheres

Conventional 2D cultures on *in vitro* substrates enable rapid drug equilibration but fail to model diffusion resistance and architectural constraints of solid tumors. Therefore, we used 3D glioblastoma spheroids generated to maintain spatially confined growth, native cell–cell interfaces, and physiological cortical actin organisation under cytotoxic challenge. The gliomaspheres cultured showed enrichment for Nestin and CD44-positive cells, which are characteristic of GBM and make them realistic representatives of GBM (**Figure S3A**). When treated with increasing concentrations of the drug, NanoEpi exhibited significantly greater accumulation in the system than free Epi (**Figure 3A-3B**). The size of the tumorspheres decreased after 24 hours of drug treatment in both the Epi and NanoEpi categories (**Figure S3B**). This improved uptake translated into increased DNA damage, as indicated by p-H2AX, and apoptotic zones, as indicated by cleaved caspase-3, throughout the spheroid at equivalent doses (**Figure 3C, S3C**). This validates that these carriers, resistant to oxidation and efficient at penetration, are especially well-suited for delivering anthracyclines over sustained periods within dense, barrier-laden 3D tumor structures (**Figure 3C, S3C**). Moreover, in the Matrigel invasion assay, both 20 µM Epi and NanoEpi treated Gli5 tumorspheres showed disintegration by Day 3, whereas 5 µM NanoEpi spheroids exhibited marked invasion suppression relative to free epirubicin, which showed minimal detectable invasion area at 5 µM (**Figure 3D**). Actin protrusion morphologies co-stained by phalloidin/DAPI confirmed the absence of matrix-infiltrating filaments in NanoEpi, suggesting a disruption of the migratory machinery required for glioblastoma infiltration (**Figure 3E, F**).

**Figure 3:**
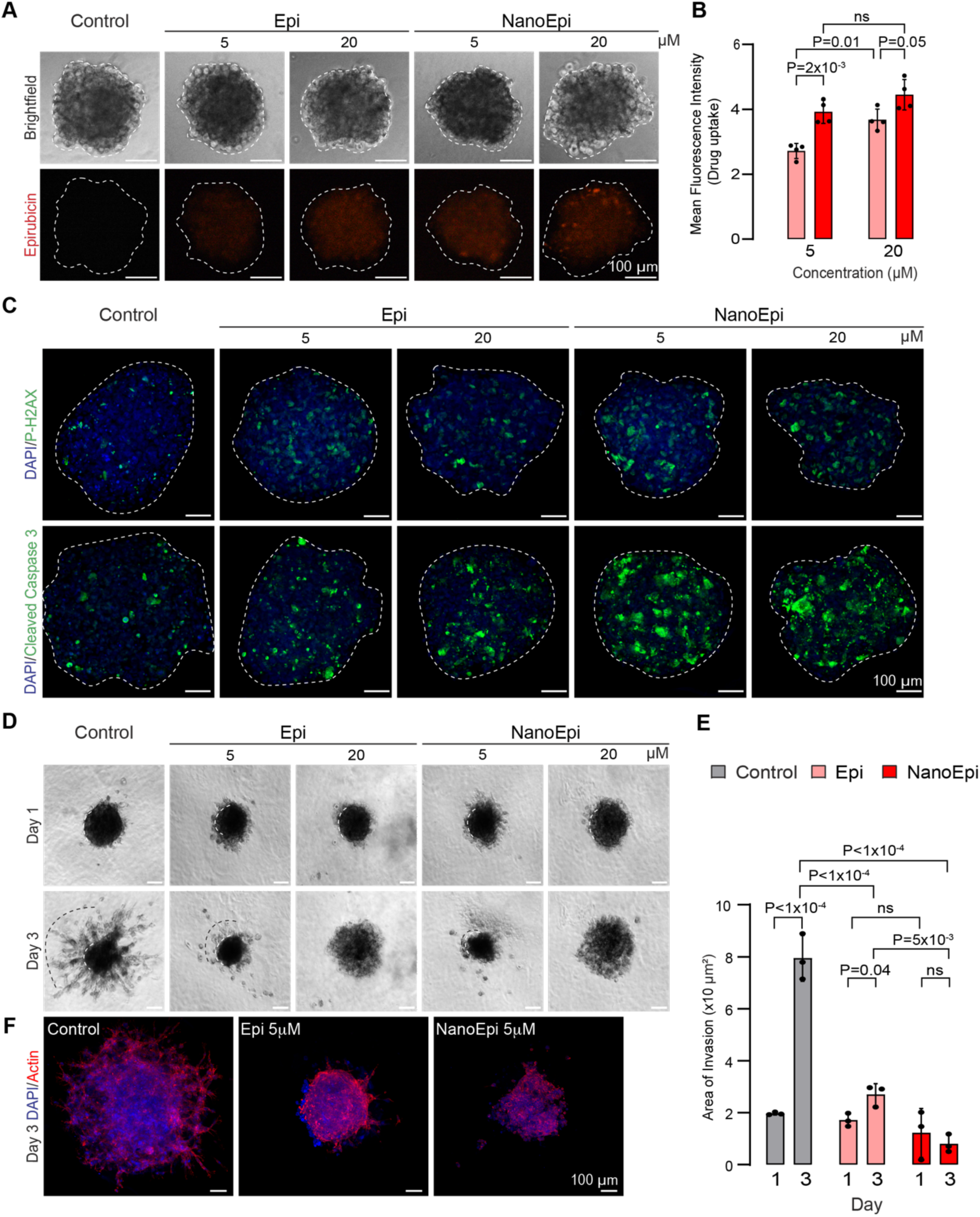
NanoEpi shows higher penetration in gliomaspheres and enhanced suppression of 3D invasion. **(A, B)** Evaluation of drug penetration in 3D gliomaspheres via fluorescence imaging, presented alongside corresponding brightfield images. Quantitative mean fluorescence intensity confirms significantly higher drug uptake with NanoEpi than with free Epi at 5 and 20 μM. **(C)** Confocal microscopy images of 3D spheres stained for DNA damage (p-H2AX, green) and apoptosis (Cleaved Caspase 3, green), demonstrating enhanced therapeutic effect in NanoEpi-treated spheres. **(D, E)** Invasion assays showing the progression of tumor cell from Day 1 to Day 3. NanoEpi treatment significantly restricts the area of invasion compared to the untreated Control and free Epi groups. (**F)** High-resolution fluorescence images (Day 3) stained for DAPI (blue) and Actin (red) reveal prominent actin-based microtentacles in the control spheres, which are drastically reduced following NanoEpi treatment.

### 3.4 NanoEpi suppresses stem-like recovery and invasive signaling and undergoes preferential lysosomal uptake

Bulk viability metrics often fail to resolve changes in sphere-initiating cancer stem-like frequency. This prompted us to apply a stringent extreme limiting dilution analysis. Using 3D tumour sphere recovery as a functional readout of the retained stem-like pool, we treated the cultures for 24 h with Nanoarch, Epi, or NanoEpi, then completely removed the drug pressure prior to assay initiation.

Following drug exposure, viable single-cell suspensions were diluted to a limiting concentration of 1 to 100 cells per well as indicated. The binary score outcome matrices were fit to a single-hit Poisson model using ELDA maximum-likelihood frequency estimation. The estimated stem cell frequency decreased from approximately 1 in 2.92 cells in the Nanoarch group to 1 in 18.23 cells (low-dose NanoEpi) and 1 in 134.92 cells (high-dose NanoEpi), representing an approximately 6.2-fold and 46.2-fold depletion of sphere-initiating cells, respectively (NA vs NAE: χ² = 18.9, *P* = 1.37 × 10⁻⁵; χ² = 62.5, *P* = 2.65 × 10⁻¹⁵). Although NanoEpi showed numerically lower stem cell frequencies than free Epirubicin at both doses (E1 (1 in 12.33 cells): NE1 (1 in 18.23 cells), E5 (1 in 56.49 cells): NE5 (1 in 134.92 cells), these differences did not reach statistical significance. These findings demonstrate that NanoEpi substantially depletes the functional stem-like compartment beyond the effect of the nanocarrier alone, supporting its superior capacity to eliminate tumour-initiating cells rather than merely delaying sphere outgrowth. (**Figure 4A**). In a parallel colony-formation experiment using the same post-treated pool, clonogenic recovery was confirmed to be lost. NanoEpi-treated cultures produced no proliferative rescue colonies, whereas free-drug spheroids retained ∼7–12% rescue at matched exposures (**Figure 4B**). To mechanistically link these phenotypic outcomes to the secretion of invasion markers, such as Matrix metalloproteinases (MMPs), 48-h conditioned supernatants collected after 24-h exposure were probed for MMP-2 by western blotting. We observed reduced MMP-2 secretion from the Epi and NanoEpi treatment groups compared to the controls (**Figure 4C**). To achieve higher sensitivity in terms of measuring MMP activity we performed gelatin zymography to quantify enzymatic function. This assay showed that NanoEpi fully eliminated the active gelatinolytic forms of MMP-2 and MMP-9 in Gli5 and U251-MG cells, correlating with the blockade of invasion (**Figure 4D**). To determine whether post-treatment proliferative rebound differed in 3D tumor architecture, we withdrew both formulations from spheroids and quantified cell cycling 48 h later by Ki-67 immunostaining. We observed a strong rebound in proliferative cells under free-drug exposure (Epi), whereas NanoEpi-treated spheroids remained largely quiescent, with negligible Ki-67 signal, indicating prolonged inhibition of proliferative recovery (**Figure 4E**). Given this prolonged suppression following transient exposure to NanoEpi, we next examined whether intracellular lysosomal retention contributed to the sustained response. Confocal imaging further demonstrated preferential lysosomal localisation of NanoEpi compared to free epirubicin, suggesting lysosome-mediated intracellular retention of NanoEpi and its sustained release within GBM cells (**Figure 4F**).

**Figure 4:**
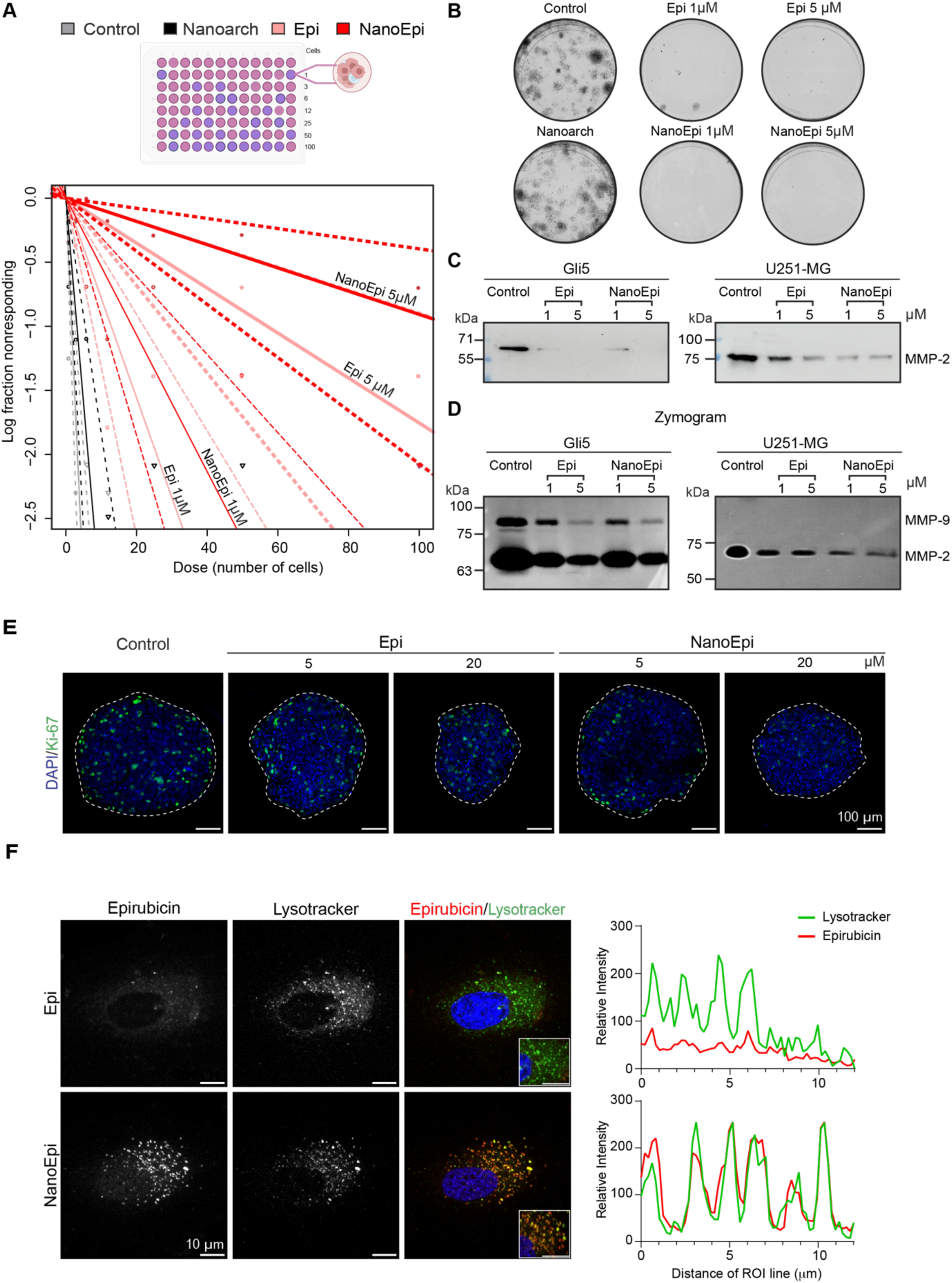
NanoEpi suppresses stem-like self-renewal and matrix-metalloproteinases in GBM. **(A)** Extreme Limiting Dilution Assay (ELDA) representing the log fraction of non-responders versus the number of cells seeded. Steeper slopes for NanoEpi-treated groups (NanoEpi1, NanoEpi5) compared to free Epirubicin (Epi1, Epi5) and controls (Con, Nanoarch) indicate reduction in the frequency of stem-like initiating cells. **(B)** Representative images of colony formation assay showing the long-term proliferative potential of GBM cells. Treatment with both free Epirubicin and NanoEpi results in a marked reduction in colony density, with NanoEpi-treated groups showing nearly complete inhibition at higher concentrations. **(C)** Western blot analysis of Matrix Metalloproteinase MMP2 in secreted fractions. **(D)** Gelatin Zymography showing that NanoEpi treatment results in a significant downregulation of MMP2 and MMP9 secretion compared to the control, correlating with a reduced invasive potential. **(E)** Immunofluorescence imaging of 3D gliomaspheres stained for DAPI (blue) and the proliferation marker Ki-67 (green). A dose-dependent decrease in Ki-67 positive cells is observed in NanoEpi-treated spheres (5 μM and 20 μM) compared to the control and free Epi groups, indicating effective suppression of tumor cell proliferation within the 3D microenvironment even after removal of the drug. **(F)** Confocal images depicting co-localization of Lysotracker and epirubicin in Epi- and NanoEpi-treated cells. NanoEpi-treated cells show enhanced lysosomal localisation, suggesting lysosome-mediated uptake and sustained intracellular drug release. The insets show a representative 3D-rendered region of the cell. The line profile on the right represents the co-localisation intensity profile.

Recently, we showed an increased cumulative drug release from loaded nanoarchaeosomes at acidic pH [16]. A PEG-based acid-labile conjugate of epirubicin enables pH-dependent release, efficiently killing cancer cells and reducing cardiotoxicity [20]. Lysosome as an organelle marks the fitness for GBM cells by regulating mTOR signalling and preserving the trafficking machinery for proper receptor accumulation in membrane [21]. Owing to their ability to regulate the signalling and degradation pathways, lysosomes are essential for glioblastoma plasticity. Therefore, hijacking the lysosomal routes for drug delivery is more efficient for better outcomes. To assess this we treated the cells with free durg or NanoEpi and performed colocalization analysis with lysotracker dye. We observed a typical punctate accumulation of NanoEpi in the lysosomes, in contrast to a more diffused Epi signal (**Figure 4F**). Thus, we propose that nanoarchaeosomes are internalised via the standard endo-lysosomal uptake route. This leads to greater epirubicin buildup within lysosomes, positioning NanoEpi closer to the nucleus, consistent with the stronger nuclear Epi signal observed compared to free Epi (**Figure 4F**).This correlates with the increased p-H2AX signal observed in the NanoEpi-treated Gli5 cells.

### 3.5 NanoEpi suppresses tumor-induced angiogenesis in the CAM model

To determine whether nanoarchaeosome-based delivery modulates tumour-driven angiogenic induction at a functional vascular interface, we performed an ex-ovo chick chorioallantoic membrane (CAM) implantation model using fertilised embryos. Peri-implant vasculature was imaged longitudinally under a stereomicroscope to map branch density and radial vessel convergence, while matched implant-proximal CAM tissue was harvested for transcript analysis. In the CAM vasculature, the tumour microenvironment (TME) implants significantly increased the number of nodes, vessel length, and area. Upon treatment with NanoEpi, we observed that these changes were significantly suppressed, demonstrating the efficacy of NanoEpi (**Figure 5A-D**). We also observed that NanoEpi treatment downregulated the transcript levels of the angiogenic drivers VEGFA, ANG1, and FGF2 relative to untreated vascular TME implants, whereas the Nanoarch vehicle exerted no significant depletion of these transcripts, confirming that bioactivity was formulation-dependent and drug-driven (**Figure 5E**).

**Figure 5.**
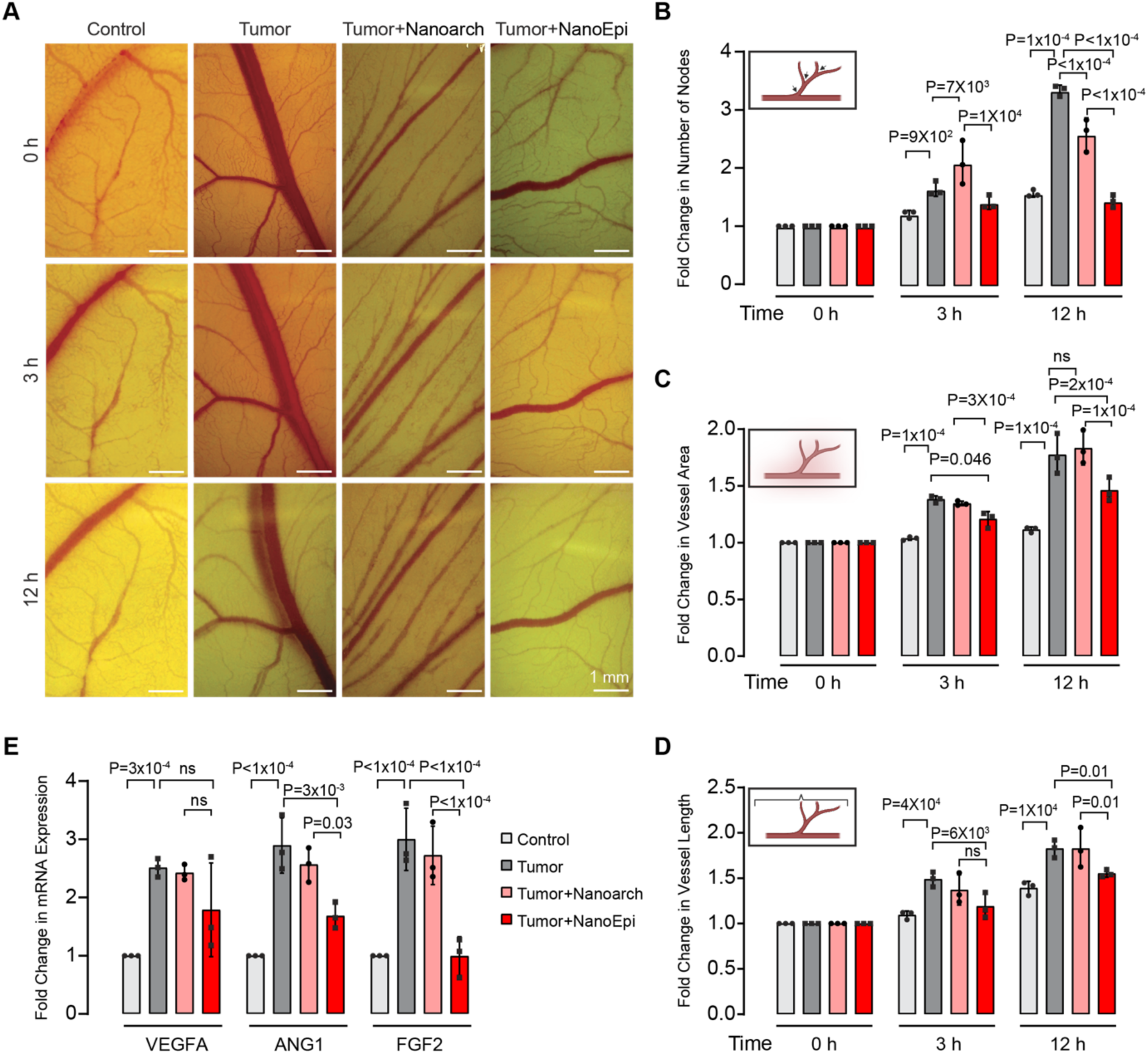
Nanoarcheosome-loaded Epirubicin suppresses tumor-induced angiogenesis and angiogenic gene expression in the CAM model. **(A)** Representative stereomicroscopic images of the chorioallantoic membrane (CAM) vasculature over a 12-hour period (0 h, 3 h, 12 h). Groups include a non-tumor Control, a tumor-only group (Gli5), tumor + Nanoarch, and tumour + NanoEpi. The NanoEpi-treated group shows a visible reduction in the density and branching of the recruitment vessels compared to the tumor-only and Nanoarch controls. **(B–D)** Quantitative morphometric analysis of the vascular NanoEpi effect over time, displaying the fold change **(B)** Number of Nodes: NanoEpi treatment significantly restricts the formation of new vascular branching points at 12 h compared to the tumor group **(C)** Vessel Area: A reduction in total vascularized area is observed in the NanoEpi group relative to the untreated tumor at the 12 h time point. **(D)** Vessel Length: NanoEpi treatment limits the overall expansion of vessel length, demonstrating superior inhibitory effects. **(E)** Quantitative RT-PCR analysis of pro-angiogenic mRNA expression in CAM tissues. Treatment with NanoEpi downregulates key angiogenic factors, including VEGFA, ANG1, and FGF2, compared to the tumor and tumour + Nanoarch groups. These results indicate that NanoEpi effectively disrupts the molecular signalling required for tumour-induced neovascularisation.

The convergence of functional sphere-initiating depletion, complete loss of clonogenic rescue, elimination of active gelatinolytic MMP-2/MMP-9 species, durable suppression of Ki-67 in 3D spheroids upon withdrawal of NanoEpi, and significant downregulation of angiogenic induction at a live vascular microinterface collectively establish the therapeutic potential of NanoEpi. Moreover, Epi-loaded nanoarchaeosome not only restricts acute cytotoxic responses but also selectively erodes the post-treatment regenerative cancer stem-like compartment, prevents protease recovery linked to invasion, blocks proliferative resurgence after drug removal, and suppresses tumour-driven neovascular engagement at spatially relevant 3D tumour-vascular micro-interfaces. These findings establish nanoarchaeosomes as a high-resilience nanocarrier platform for future anthracycline -delivery translation in GBM.

## 4. Conclusions

Glioblastoma continues to resist durable therapeutic control due to its invasive and stem-like tumor cell populations. In this study, we demonstrate that delivering epirubicin through or via nanoarchaeosome encapsulation enhances the intracellular stress signaling while suppressing the adaptive tumor phenotypes associated with recurrence..

NanoEpi induced significantly more DNA damage than free epirubicin, despite similar cellular uptake, indicating enhanced intracellular retention and drug engagement. These findings are consistent with the observed mitochondrial dysfunction and ROS accumulation. Notably, NanoEpi preferentially accumulated within lysosomal compartments, suggesting that this lysosome-associated sequestration may contribute to sustained intracellular release and prolonged stress signaling. In patient-derived 3D gliomaspheres, NanoEpi showed enhanced penetration and retention, which in turn suppressed invasion and disrupted actin-rich protrusive structures linked to glioblastoma infiltration.

NanoEpi also reduced MMP activity and tumour-induced angiogenesis in the CAM model, indicating a wider impact on invasion-related and microenvironmental signalling pathways.

Collectively, these findings support a model in which nanoarchaeosome-mediated lysosomal retention enhances intracellular epirubicin activity and suppresses glioblastoma’s adaptive survival programs. In addition to improving drug delivery efficiency, nanoarchaeosomes actively reshape tumor responses to cytotoxic stress, underscoring their potential as a translational platform for GBM therapy. With the advent of newer therapies targeting specific signalling pathways, our findings with Nanoarchaeosomes can provide an enhanced therapeutic opportunity. Future work aimed at identifying the precise pathways affected by these nanoarchaeosome formulations could reveal new vulnerabilities in glioma cells to target, further improving treatment outcomes. Considering the expanding range of therapies designed to target specific signalling pathways, our findings suggest that Nanoarcheosomes could broaden opportunities for effective therapeutic interventions, with significant implications for public health and for realising precision medicine.

## Author Contributions

A.S.G., S.S. and P.L.C. conceived the original idea. A.S.G. and S.A. synthesised and characterised the nanoarchaeosomes. S.G. and S.M. generated patient cell lines and characterised them. A.S.G., S.G, A.H, and P.L.C. performed the formal data analysis. A.S.G. and P.L.C. wrote the original manuscript draft. S.S., S.G., and S.A. contributed to the discussion, revision, and completion of the manuscript. S.S. and P.L.C. supervised the work. All authors contributed to the scientific discussion and revision of the manuscript.

## Notes

The authors declare no competing financial interest.

## Acknowledgments

This work was supported by the Wellcome Trust/DBT India Alliance [IA/I/19/1/504280 to PLC], SERB Power Grant [SPG/2022/001987]; UGC [Fellowship No. 231620157176 A.S.]; and IISER Tirupati core funds. The authors also thank IIT Madras and the Government of India for financial assistance through the following grants: SERB [SP23242582AMSERB009000]; Ministry of Earth Sciences [SP24250515CHMOES008504]; INAE [SP24251232AMINAE009000]; and DBT [SP25261072AMDBTX009000]. We thank Dr S. Chavali, IISER Tirupati, for useful discussions and comments.

## References

1. Pouyan, A., et al., Glioblastoma multiforme: insights into pathogenesis, key signaling pathways, and therapeutic strategies. Mol Cancer, 2025. 24(1): p. 58.

2. Alves, A.L.V., et al., Role of glioblastoma stem cells in cancer therapeutic resistance: a perspective on antineoplastic agents from natural sources and chemical derivatives. Stem Cell Res Ther, 2021. 12(1): p. 206.

3. Yang, F., et al., Delivery of epirubicin via slow infusion as a strategy to mitigate chemotherapy-induced cardiotoxicity. PLoS One, 2017. 12(11): p. e0188025.

4. Fabel, K., et al., Long-term stabilization in patients with malignant glioma after treatment with liposomal doxorubicin. Cancer, 2001. 92(7): p. 1936–42.

5. Han, Y. and J.H. Park, Convection-enhanced delivery of liposomal drugs for effective treatment of glioblastoma multiforme. Drug Deliv Transl Res, 2020. 10(6): p. 1876–1887.

6. Jahangiri, A., et al., Convection-enhanced delivery in glioblastoma: a review of preclinical and clinical studies. J Neurosurg, 2017. 126(1): p. 191–200.

7. Vanbilloen, W.J.F., et al., Nanoparticle Strategies to Improve the Delivery of Anticancer Drugs across the Blood-Brain Barrier to Treat Brain Tumors. Pharmaceutics, 2023. 15(7).

8. Parvar, S.J., et al., Convection-enhanced delivery for brain malignancies: Technical parameters, formulation strategies and clinical perspectives. Adv Drug Deliv Rev, 2025. 224: p. 115657.

9. Zhu, D., et al., Enhancing TIM-3 immunotherapy with epirubicin-loaded pH sensitive fusion membrane nanoparticles for effective glioblastoma treatment. J Control Release, 2025. 388(Pt 2): p. 114367.

10. Patel, G.B. and G.D. Sprott, Archaeobacterial ether lipid liposomes (archaeosomes) as novel vaccine and drug delivery systems. Crit Rev Biotechnol, 1999. 19(4): p. 317–57.

11. Kaur, G., et al., Archaeosomes: an excellent carrier for drug and cell delivery. Drug Deliv, 2016. 23(7): p. 2497–2512.

12. Patel, G.B., et al., In vitro assessment of archaeosome stability for developing oral delivery systems. Int J Pharm, 2000. 194(1): p. 39–49.

13. Benvegnu, T., L. Lemiegre, and S. Cammas-Marion, New generation of liposomes called archaeosomes based on natural or synthetic archaeal lipids as innovative formulations for drug delivery. Recent Pat Drug Deliv Formul, 2009. 3(3): p. 206–20.

14. Babunagappan, K.V., et al., Doxorubicin loaded thermostable nanoarchaeosomes: a next-generation drug carrier for breast cancer therapeutics. Nanoscale Adv, 2024. 6(8): p. 2026–2037.

15. Basha, S.S., et al., Nanoarchaeosomes for synergistic photochemotherapy in triple-negative breast cancer. Sci Rep, 2025. 16(1): p. 460.

16. Ariraman, S., et al., Paclitaxel-Loaded Nanoarchaeosomes: An Innovative Strategy for Targeted Breast Cancer Treatment. Langmuir, 2026. 42(5): p. 3937–3949.

17. Hu, Y. and G.K. Smyth, ELDA: extreme limiting dilution analysis for comparing depleted and enriched populations in stem cell and other assays. J Immunol Methods, 2009. 347(1-2): p. 70–8.

18. Cersosimo, R.J. and W.K. Hong, Epirubicin: a review of the pharmacology, clinical activity, and adverse effects of an adriamycin analogue. J Clin Oncol, 1986. 4(3): p. 425–39.

19. Ong, M.S., et al., Cytoskeletal Proteins in Cancer and Intracellular Stress: A Therapeutic Perspective. Cancers (Basel), 2020. 12(1).

20. Takahashi, A., et al., NC-6300, an epirubicin-incorporating micelle, extends the antitumor effect and reduces the cardiotoxicity of epirubicin. Cancer Sci, 2013. 104(7): p. 920–5.

21. Merlet, L., M. Le Guyon, and J. Gavard, A lysosomal requiem for glioblastoma cells. Trends Cancer, 2026. 12(5): p. 386–389.

