## Supplementary figure for "Nanoarchaeosome-mediated epirubicin delivery induces sustained intracellular stress and suppresses adaptive glioblastoma phentoypes"

### Supplementary Figures

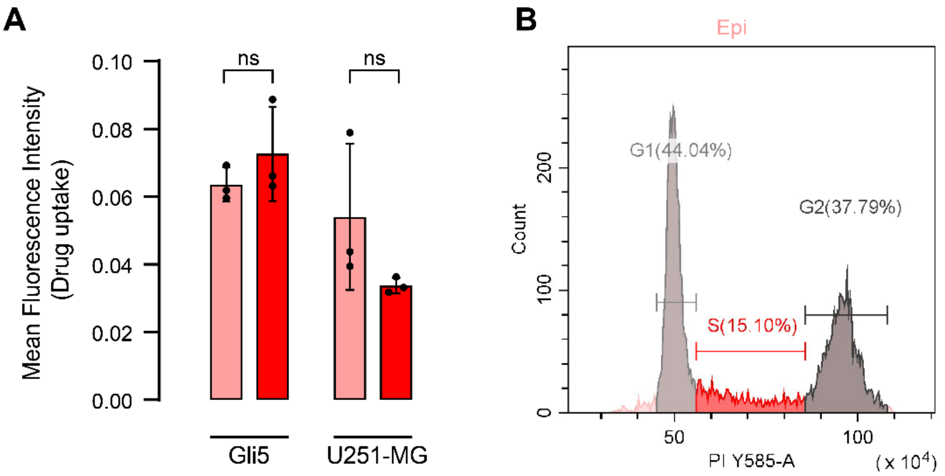


**Figure S1:** (A) The mean fluorescence intensity of drug uptake in monolayer cells (Gli5 and U251-MG) treated with free Epirubicin and NanoEpi shows no significant difference in the drug uptake. (B) The cell cycle profile of Gli5 cells treated with free Epi also shows S-phase arrest supporting the similar activity after loading in nanoarchaeosomes.

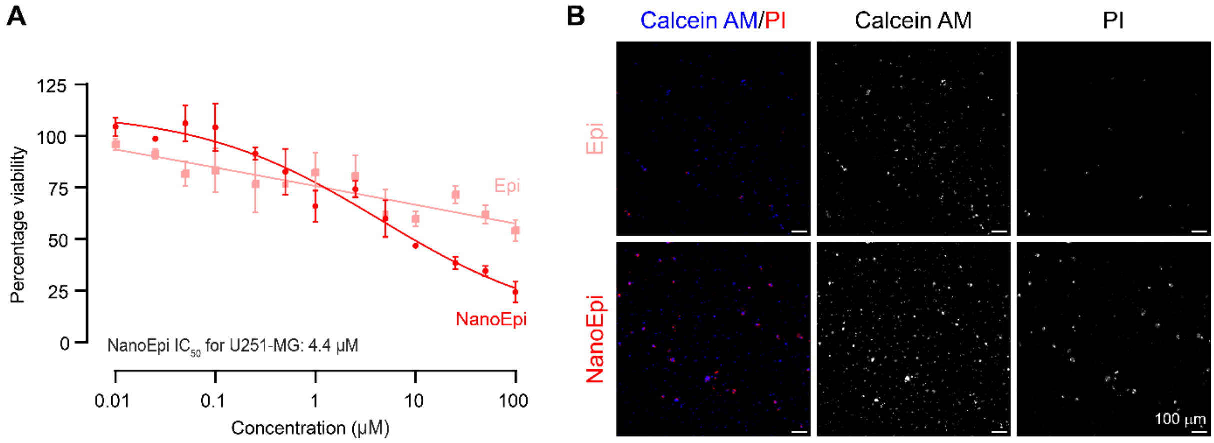


**Figure S2:** (A) The $IC_{50}$ curve generated for Epi and NanoEpi when treated to the U251-MG cells indicating the better efficacy of NanoEpi than free Epi with a $IC_{50}$ value of 4.4 µM. (B) The live/dead staining of Gli5 cells treated with 5 µM Epi and NanoEpi showing a proportionate increase in the dead cells marked by Propidium Iodide (2 µg/mL, Merck) penetration comparing the live cells fluorescing with Calcein AM (5 µM, Invitrogen).

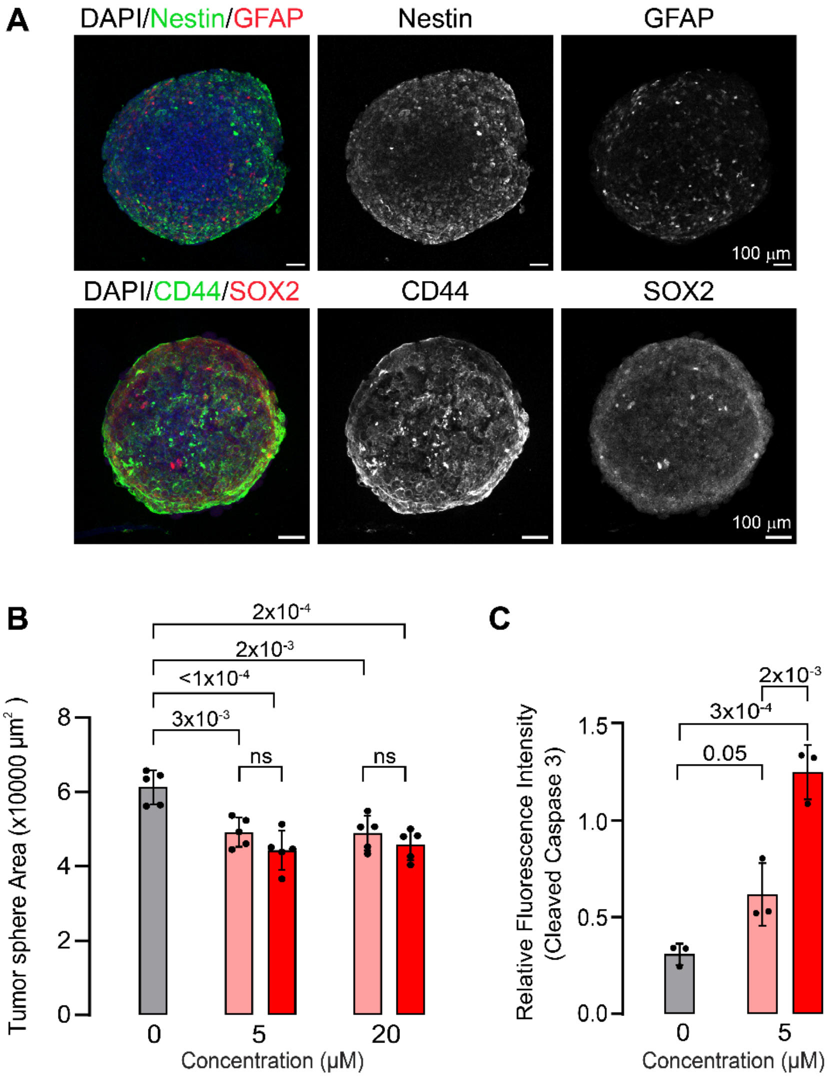


**Figure S3:** (A) A solid population of cells in Gli5 gliomaspheres expresses Nestin and CD44 making the use of this 3D model a more realistic disease state relevant one. (B) The tumor sphere area measured after 24 hours of drug treatment shows reduced area after both Epi and NanoEpi treatment at 5 and 20 µM concentrations. (C) Quantification of fluorescence intensity of Cleaved Caspase-3, after 5 µM treatment of Epi and NanoEpi shows that NanoEpi significantly induced more apoptosis throughout the 3D spheroid.

The primary antibodies used includes Human Nestin (1:300, MAB1259, R&D Systems), CD44 (IM7) (1:500, 14-0441-82, Invitrogen), GFAP (D1F4Q) (1:300, 12389, CST) and SOX2 (1:300, ab97959, Abcam).
